# Effects of chondrogenic priming duration on mechanoregulation of engineered cartilage anlagen

**DOI:** 10.1101/2020.09.02.280115

**Authors:** Anna M. McDermott, Emily A. Eastburn, Daniel J. Kelly, Joel D. Boerckel

## Abstract

Bone development and repair occur by endochondral ossification of a cartilage anlage, or template. Endochondral ossification is regulated by mechanical cues. Recently, we found that in vivo mechanical loading promoted regeneration of large bone defects through endochondral ossification, in a manner dependent on the timing of load initiation. Here, we have developed an in vitro model of the cartilage anlage to test whether the chondrogenic differentiation state alters the response to dynamic mechanical compression. We cultured human bone marrow stromal cells (hMSCs) at high cell density in fibrin hydrogels under chondrogenic priming conditions for periods of 0, 2, 4, or 6 weeks prior to two weeks of dynamic mechanical loading. Samples were evaluated by biomechanical testing, biochemical analysis of collagen and glycosaminoglycan (GAG) deposition, gene expression analysis, and immunohistological analysis, in comparison to time-matched controls cultured under static conditions. We found that dynamic loading increased the mechanical stiffness of engineered anlagen in a manner dependent on the duration of chondrogenic priming prior to load initiation. For chondrogenic priming times of 2 weeks or greater, dynamic loading enhanced the expression of type II collagen and aggrecan, although no significant changes in overall levels of matrix deposition was observed. For priming periods less than 4 weeks, dynamic loading generally supressed markers of hypertrophy and osteogenesis, although this was not observed if the priming period was extended to 6 weeks, where loading instead enhanced the expression of type X collagen. Taken together, these data demonstrate that the duration of chondrogenic priming regulates the endochondral response to dynamic mechanical compression in vitro, which may contribute to the effects of mechanical loading on endochondral bone development, repair, and regeneration in vivo.

## Introduction

Endochondral ossification is the dominant mode of bone formation during development and natural bone fracture repair [1], [2]. Endochondral ossification initiates by formation of a cartilage anlage, or template, which undergoes chondrocyte maturation and hypertrophy. Then, neovascular invasion results in remodeling of the hypertrophic cartilage and enables osteoblastogenesis and bone formation. The power of the endochondral program to promote rapid bone formation during development and natural repair has inspired tissue engineers to recapitulate this process for the regeneration of large bone defects, which cannot heal on their own, and do not naturally undergo endochondral ossification [3]–[9].

Mechanical forces direct endochondral bone formation. In 1881 Wilhelm Roux developed the concept of “Entwicklungsmechanik” or developmental mechanics, in which he proposed that different mechanical forces in development give rise to different tissues (e.g., compression promotes bone formation) [10]. Mechanical forces exerted in utero are critical for proper development of the human skeleton[11]–[14]. Similarly, for endochondral bone repair, foundational studies from Stephan Perren [15], [16], and others [17]–[23], demonstrate that mechanical cues, induced by interfragmentary motion in the fracture gap, directly determine tissue differentiation of the callus. Based on these observations, we hypothesized that mechanical forces would be similarly important for regulating the process by which cartilage is converted into bone. Using a development-mimetic approach, we found that mechanical loading significantly enhanced endochondral bone defect regeneration in vivo [5], [24]. Importantly, the endochondral regeneration response to loading depended on the timing of load initiation [5]. Early loading, applied continuously, prolonged cartilage persistence and impaired vascular ingrowth, while delayed loading, initiated after the onset of chondrocyte hypertrophy, accelerated endochondral ossification and enhanced neovascularization [5], [25].

Because endochondral ossification requires a concert of cell types (i.e., chondrocytes, endothelial cells, and osteoblasts), this raises the question of which of these cell types are the critical mechano-responders and whether the effect of the timing of load initiation on endochondral ossification is autonomous to a given cell type or is due to their coordinated interactions. We set out to address this question by engineering each of these tissues in vitro, to directly test the effects of load timing on each tissue in isolation. Using engineered neovascular networks in vitro, we recently found that early loading directly disrupted neovessel formation, while delayed loading promoted neovascular mechanotransduction and angiogenesis [26]. Similarly, using engineered cartilage anlagen in vitro, we and others found that both early and delayed loading promoted chondrogenesis [5], [27]–[30] with delayed loading enhancing production of the angiogenic growth factor, VEGF [5], which both shows that chondrocytes are autonomously mechanoresponsive and may suggest a role for loading in directing non-autonomous chondrocyte-endothelial cell crosstalk. However, the interaction between the chondrocytic differentiation/maturation state and the timing of mechanical load initiation is poorly understood. Therefore, the goal of this study was to examine the effect of chondrogenic priming duration on the cell and tissue response to dynamic compression.

Here, we chondrogenically primed fibrin hydrogel-encapsulated human bone marrow stromal cells (MSCs) to various stages of chondrogenic maturation (0, 2, 4, or 6 weeks of chondrogenic priming) prior to applying dynamic compression for two additional weeks. We then evaluated the effects of dynamic loading on tissue mechanical properties, cellular gene expression, and matrix deposition. We found that dynamic loading induced tissue stiffening and cellular mechanotransduction in manner dependent on priming duration, and suppressed mineral deposition at latter stages of differentiation, but did not significantly alter the bulk cartilage matrix composition at any time point. Together these data suggest that chondrocyte mechanotransduction depends on differentiation state and may particularly influence the transition from chondrocyte hypertrophy to ossification.

## Materials and Methods

### Study design

The goal of this study was to determine how the extent of chondrocyte maturation influences the mechanotransductive and chondrogenic response of hydrogel-embedded hMSCs to dynamic mechanical loading. We used a custom-made bioreactor to apply dynamic, unconfined compression to hMSC-laden hydrogels (**Fig. 1A,B**), which were then cultured under free-swelling conditions in chondrogenic media for either 0, 2, 4, or 6 weeks prior to 2 weeks of dynamic compression (**Fig. 1C**). Dynamically loaded samples were compared to time-matched free swelling (FS) controls. The hydrogels were collected at the end of their loading cycle for analysis, as described below.

**Figure 1:**
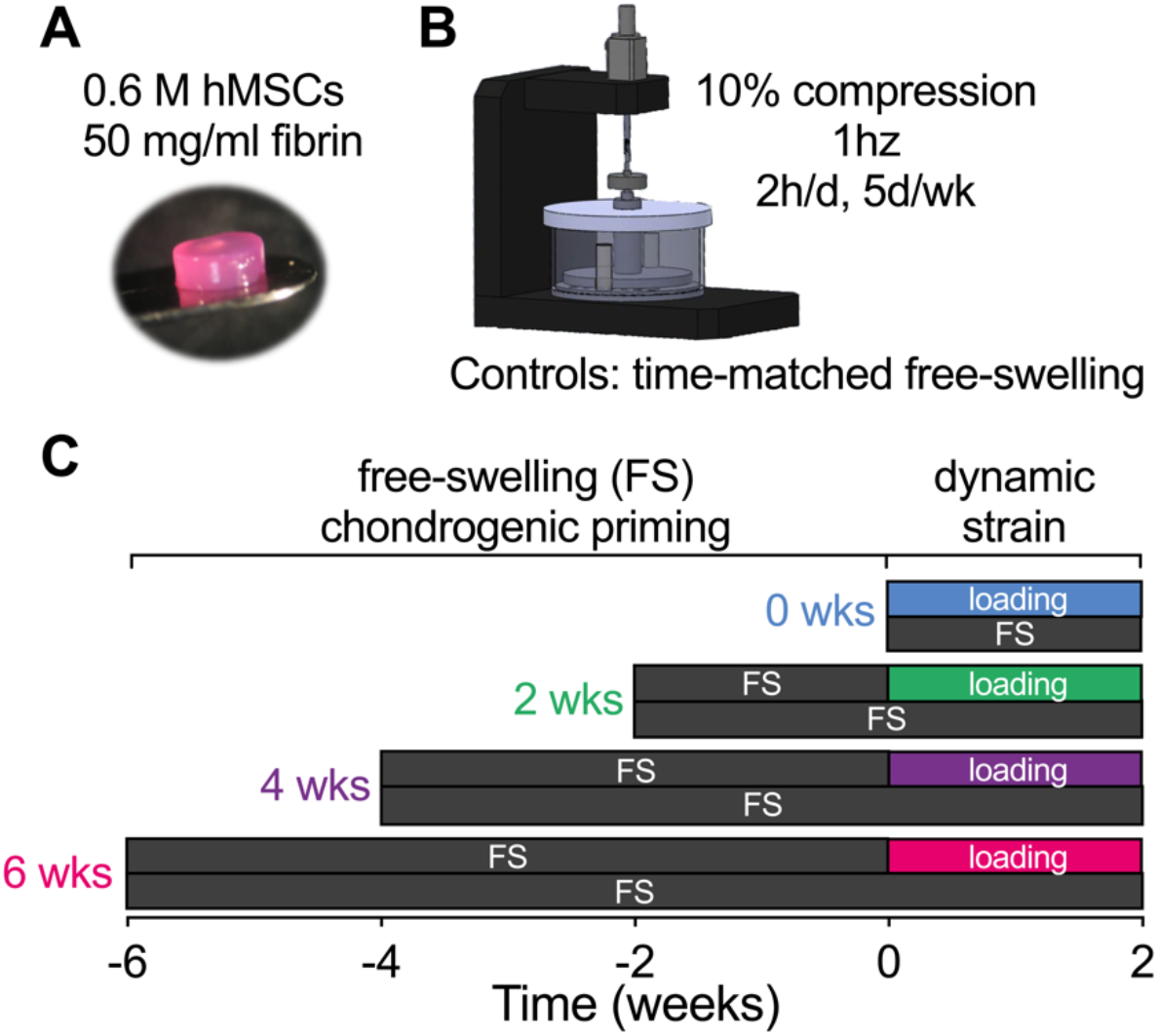
Bioreactor setup and loading timeline. (A) hMSC-laden hydrogels (P3, 589,000 cells, 50mg/mL fibrinogen) were subjected to different degrees of chondrogenic priming (0wks, 2wks, 4wks, 6wks) followed by 2 weeks of (B) dynamic compression in a custom-made bioreactor. (C) All samples were collected after their loading cycle and compared to free-swelling controls at the same time.

### Cell culture

Human bone marrow stromal cells (hMSCs, P3) (Lonza) were expanded in high glucose Dulbecco’s Modified Eagle’s Medium (4.5 mg/mL glucose, 200 mM l-glutamine, hgDMEM), supplemented with 10% fetal bovine serum (FBS), 1% Penstrep, and 5 ng/mL FGF-2. The cultures were expanded to passage 3 (P3) and maintained in a humidified environment at 37°C, 5% CO2, and 5% O2.

### Fibrin gel preparation and chondrogenic culture

Cells were then polymerized into fibrin hydrogels at 15×10^6^ cells/mL. At 80% confluency, P3 hMSCs were trypsinized and resuspended in a 10,000 KIU/mL aprotinin solution (Nordic Pharma) with 19 mg/mL sodium chloride and 100mg/mL bovine fibrinogen (Sigma-Aldrich). Fibrinogen was polymerized into fibrin by combining cell suspensions 1:1 with a solution of 5 U/mL thrombin and 40 mM CaCl2 for a final solution of 50 mg/mL fibrinogen, 2.5 U/mL thrombin, 5000 KIU/mL aprotinin, 17 mg/mL sodium chloride, 20 mM CaCl2, and 15×10^6^ cells/mL. The cell-laded hydrogel solution was then pipetted into 5 mm diameter x 2 mm thickness cylindrical agarose molds to create uniform constructs containing approximately 589,000 cells each.

Throughout the study, culture was maintained in chondrogenic media consisting of hgDMEM supplemented with 1% Penstrep, 100 KIU/mL aprotinin, 100 μg/mL sodium pyruvate, 40 μg/mL l-proline, 1.5 mg/mL bovine serum albumin, 4.7 μg/mL linoleic acid, 1x insulin-transferrin-selenium, 50 μg/mL l-ascorbic acid-2-phosphate, 100 nM dexamethasone (all Sigma-aldrich), and 10 ng/mL TGF-β3 (ProSpec-Tany TechnoGene Ltd., Israel). Fresh media was supplied every 3 days and culture was maintained in a humidified environment at 37°C, 5% CO2, and 5% O2.

### Dynamic compression

Dynamic unconfined compressive loading was applied to the constructs using a custom-made bioreactor (**Fig. 1B**). Samples were loaded under sinusoidal displacement control, applied 2 hours per day, 5 days per week at 1 Hz to an amplitude of 10% strain, after a 0.01 N preload was applied. Control was programmed using an in-house MATLAB code.

### Biochemical analysis

All constructs were harvested and analyzed at the end of their final loading cycle for DNA, sulfated glycosaminoglycan (sGAG) and collagen content (N = 5/group). Samples were digested overnight at 60°C in a solution of 125 μg/mL papain, 0.1 M sodium acetate, 5 mM ʟ-cysteine, 0.05 M EDTA (all Sigma-Aldrich). DNA content was quantified with Hoechst Bisbenzimide 33258 dye assay (Sigma-Aldrich) as described previously [31], and sulfated glycosaminoglycan content was quantified using the dimethylmethylene blue dye-binding assay (Blyscan; Biocolor Ltd.). Collagen content was quantified by measurement of orthohydroxyproline using the dimethylaminobenzaldehyde and chloramine T assay [32]. A hydroxyproline to collagen ratio of 1:7.69 was used [33] to infer total collagen content.

### RNA isolation and qPCR

Total mRNA was extracted from fibrin/hMSC constructs after the final loading cycle using RNeasy mini kit (Qiagen). RNA (300ng) was reverse transcribed using into cDNA using High Capacity cDNA Reverse Transcription Kit (Applied Biosystems). Gene expression changes relative to free swelling controls were quantified via real-time reverse transcription-polymerase chain reaction (qRT-PCR). Reactions were carried out in triplicate 20 μL volumes of 10 μL Sybr Green Master Mix (Applied Biosystems), 30 ng cDNA, 400 nM Sigma Kicqstart forward and reverse primers, on an ABI 7500 real-time PCR system (Applied Biosystems) with a profile of 95 °C for 10 min, and 40 cycles of denaturation at 95 °C for 15 sec, and annealing/amplification at 60 °C for 1 min. Quantification of target genes (Table 2) was determined against housekeeping reference gene GAPDH as fold change over free swelling controls using the delta-delta Ct method [34].

**Table 2:**
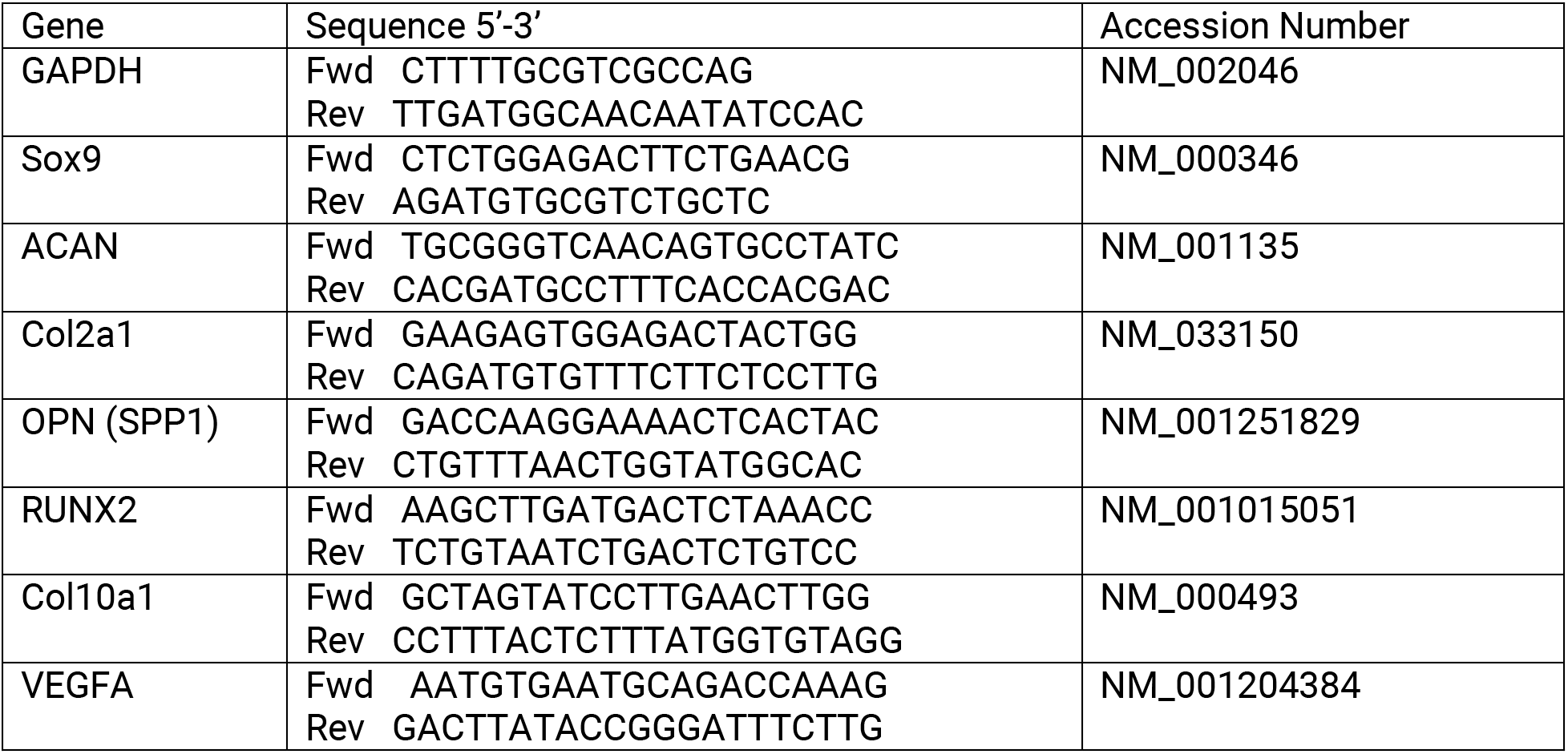
qPCR target genes

### Histology and immunohistochemistry

Hydrogels were fixed in 4% paraformaldehyde (n = 2) overnight at 4°C. Constructs were halved and paraffin embedded cut surface down and sectioned at 5 μm to provide a cross-section of the hydrogel center. Sections were cleared in xylene, rehydrated in graded alcohols, and stained for sGAG in 1% Alcian blue with 0.1% Nuclear Fast Red counter stain.

Collagen deposition was identified through immunohistochemistry (Santa Cruz). Antigen retrieval was enzyme mediated (chondroitinase ABC, Sigma-Aldrich). Slides were blocked with Innovex Background Buster and incubated with primary antibody, diluted in PBS, overnight at 4°C. Slides were washed with PBS and incubated for 10 mins with universal rabbit IgG secondary antibody, followed by 10 mins with HRP enzyme, and DAB substrate for 5 mins (all Innovex). Sections were counterstained in Hematoxylin (VWR) and mounted with Cytoseal XYL.

### Mechanical testing

Hydrogels were mechanically tested in unconfined compression (Zwick Roell Z005, Herefordshire, UK) between two steel plattens and a 5N load cell. Hydrogels were hydrated in a PBS bath at room temperature. Equilibrium modulus was obtained via a stress relaxation test where a 10% strain was applied and maintained until equilibrium was reached. Dynamic modulus was determined after 10% strain was applied for 10 cycles at 1Hz and at 0.1Hz. In both tests a preload of 0.01N was applied to ensure contact between the hydrogel and machine platens.

### Statistics

Statistical analyses were performed using either one- or two-way analysis of variance (ANOVA), as appropriate. Multiple comparisons between groups for one-way ANOVA were assessed by Tukey’s multiple comparison test and two-way by Sidak’s multiple comparison test. When necessary, data were log-transformed to ensure normality and homoscedasticity before ANOVA. Normality of dependent variables and residuals was verified by D’Agostino-Pearson omnibus and Brown-Forsythe tests, respectively. Statistical significance was set at α = 0.05.

## Results

### Mechanical properties

First, we sought to determine how construct mechanical properties are impacted by the duration of chondrogenic priming and dynamic mechanical loading. To this end, we performed mechanical testing in unconfined compression following completion of the final loading cycle for each group, in comparison to time-matched free-swelling controls (**Fig. 2**). Both equilibrium and dynamic modulus increased significantly with the duration of priming time, and dynamic loading significantly increased the equilibrium modulus under conditions of 4 weeks of priming and dynamic modulus after both 4 and 6 weeks of priming.

**Figure 2:**
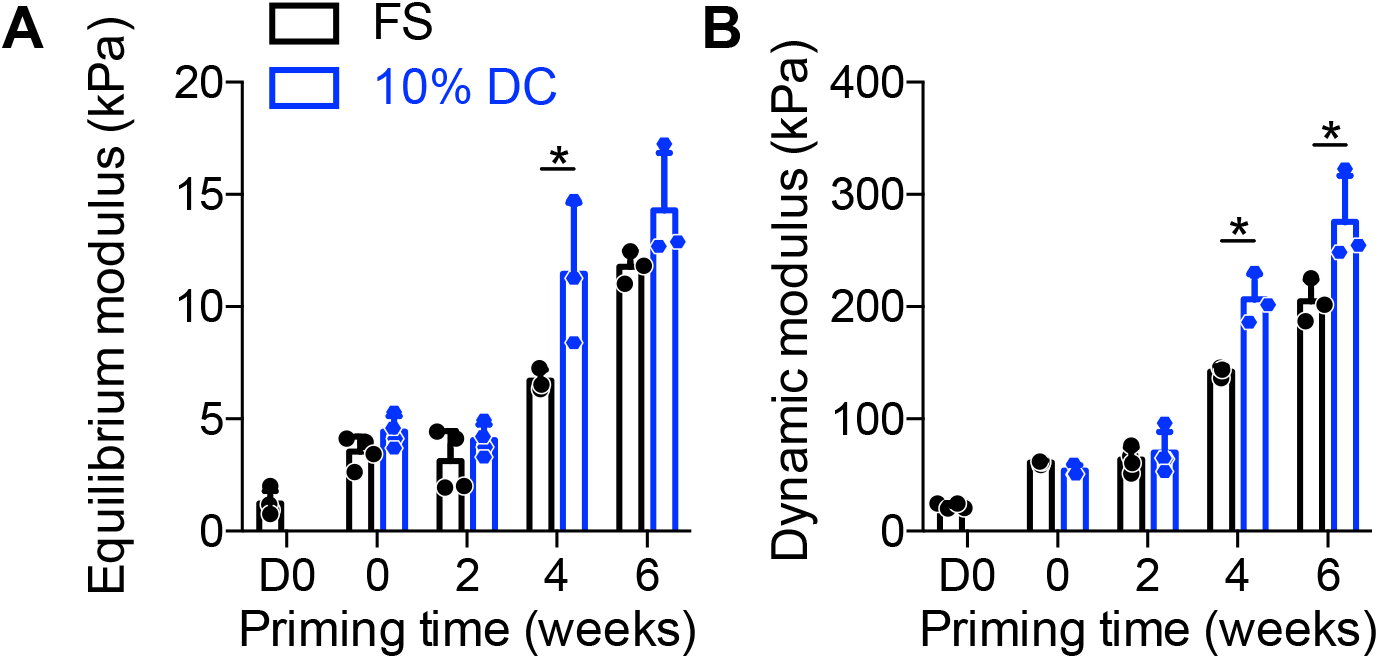
Mechanical properties of chondrogenically-primed hMSC constructs under free-swelling (FS) or 10% dynamic compression (DC) conditions. Samples were tested in unconfined compression to determine equilibrium modulus, E’ (A) and dynamic modulus, E’’ (B) at the end of the final loading cycle for each group, in comparison to time-matched free swelling controls. Data are shown as displayed as mean ± s.d. with individual data points. Differences between groups and time points were evaluated by two-way ANOVA followed by Sidak’s multiple comparison test. The significance threshold was set at α =0.05. Both equilibrium and dynamic modulus increased significantly with time. Significant differences between free-swelling (FS) and dynamic compression (DC) groups are indicated with asterisks. N = 3 – 4 per group.

### Matrix composition

To determine the means by which dynamic compression increased construct mechanical properties, we first quantified the cellular and extracellular matrix content of the constructs (**Fig. 3**). The cellular number, measured by DNA content, was increased with increasing chondrogenic priming time, but was not significantly altered by loading (**Fig. 3A**). Chondrogenic priming significantly increased glycosaminoglycan (sGAG) deposition and sGAG/DNA, but differences between loading and free-swelling groups at each time point were not statistically significant (**Fig. 3B, D**). Collagen deposition, measured by hydroxyproline content, increased with priming time but was not significantly affected by loading either total or on a per-cell basis (**Fig. 3C, E**). Histological analysis of Alcian blue-stained tissue sections revealed largely uniform acidic polysaccharide (i.e., sGAG) distribution throughout the constructs regardless of loading (**Fig. 4**). Therefore, while dynamic compression modestly influenced ECM composition overall, these data suggest that other causes are required to fully explain the load-induced changes in construct mechanical properties.

**Figure 3:**
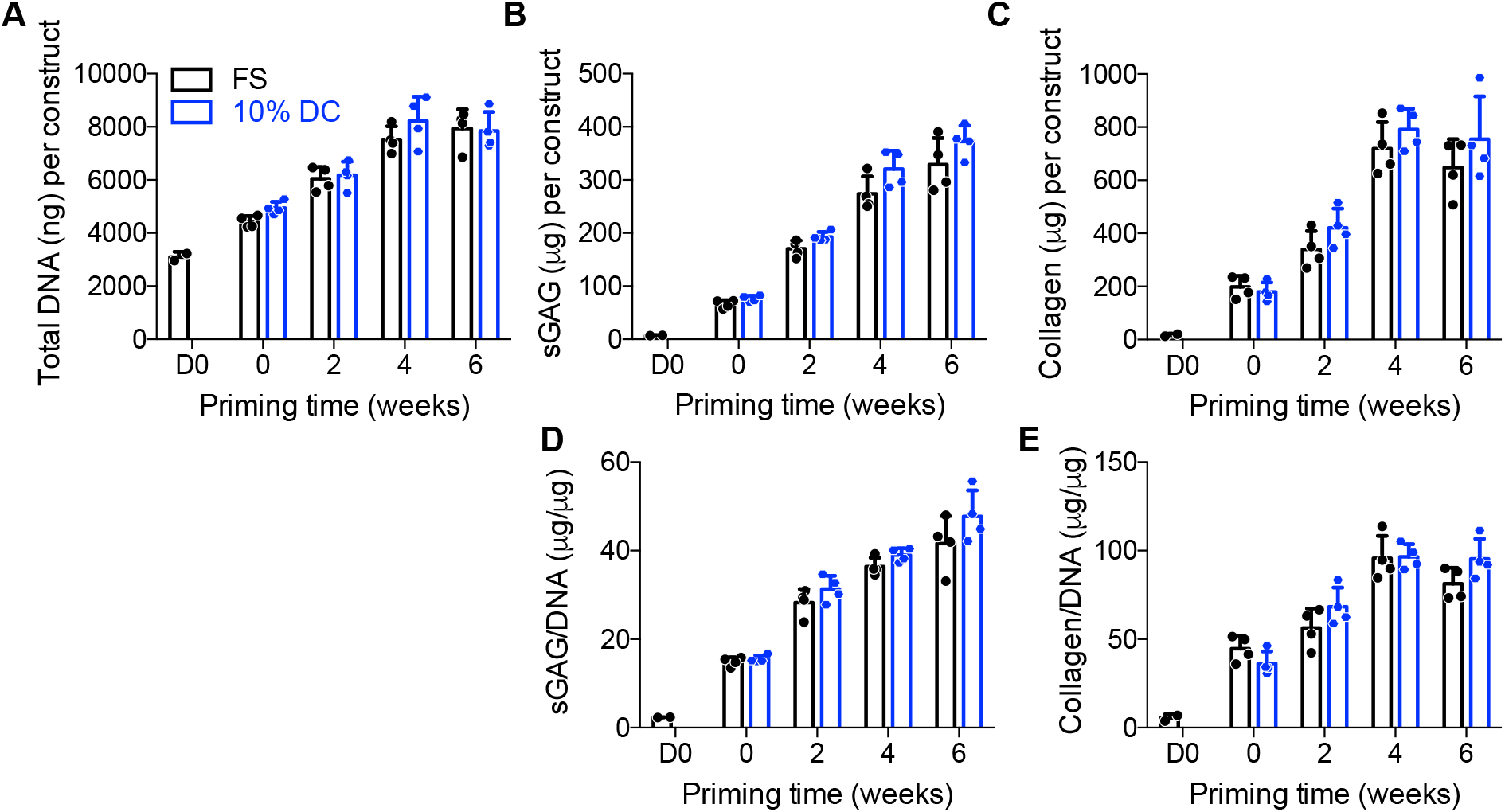
Biochemical content of chondrogenically-primed hMSC constructs under free-swelling (FS) or 10% dynamic compression (DC) conditions. Samples were digested and assayed for biochemical content at the end of the final loading cycle for each group, in comparison to time-matched free swelling controls. Each construct was assayed for total DNA content (A), total sulfated glycosaminoglycans (sGAG) (B), and total collagen (C). sGAG (D) and collagen (E) were then normalized, on a per-sample basis, to DNA content. Data are displayed as mean ± s.d. with individual data points. Differences between groups and time points were evaluated by two-way ANOVA followed by Sidak’s multiple comparison test. The significance threshold was set at α =0.05. All measures increased significantly with time. ANOVA omnibus analysis indicated a significant effect of both loading and time for sGAG and sGAG/DNA, but there were no differences for individual comparisons between FS and DC at any time point. N = 3 – 4 per group.

**Figure 4:**
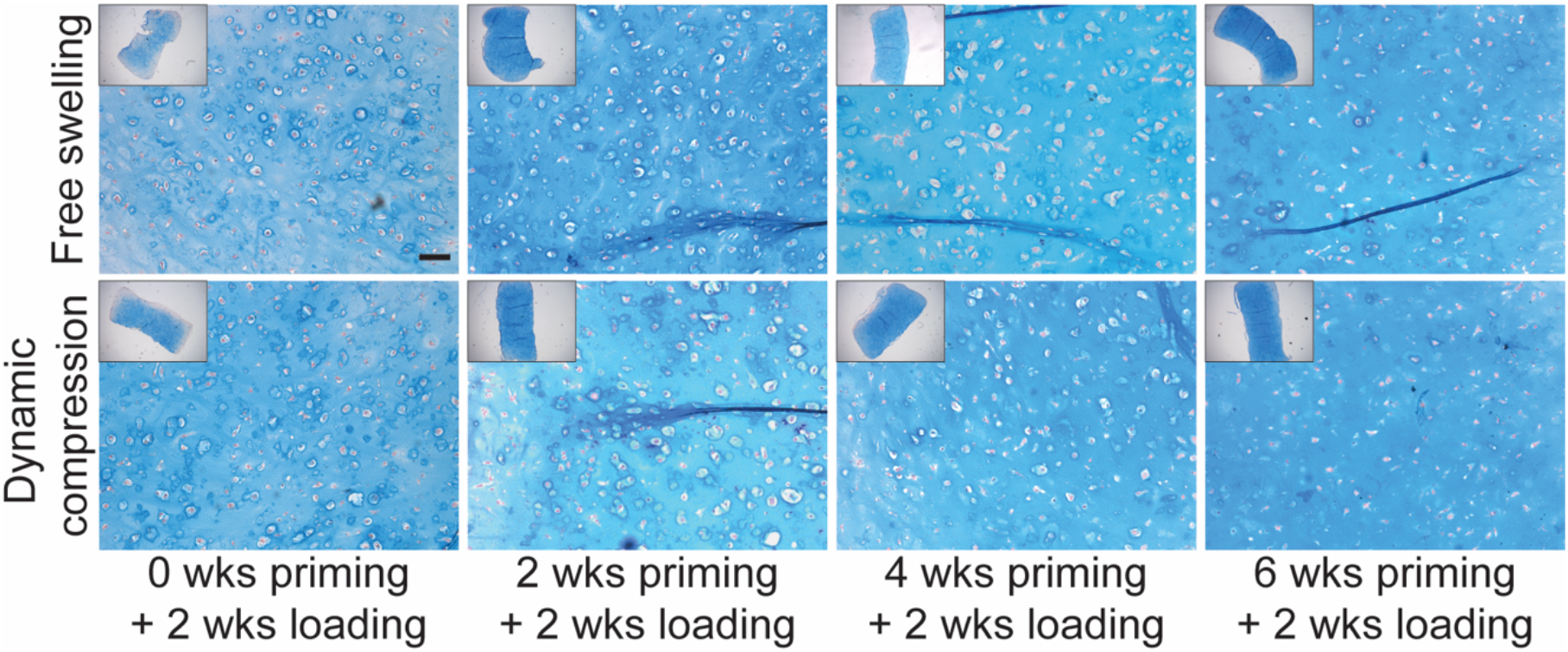
Alcian blue staining of glycosaminoglycans in chondrogenically-primed hMSC constructs under free-swelling (FS) or 10% dynamic compression (DC) conditions. Samples were fixed and sectioned at the end of the final loading cycle for each group and stained with Alcian blue/Nuclear Fast Red. Scale bar = 100 μm.

Next, to determine whether the duration of chondrogenic priming influenced transient mechanotransductive responses to dynamic compression, we evaluated load-induced messenger RNA expression of genes associated with chondrogenesis, hypertrophy, and osteogenesis, immediately after the final loading cycle in each group. We compared loaded vs. free-swelling controls at each priming time point.

### Chondrogenic gene expression

To evaluate chondrogenic gene expression, we quantified SOX9, aggrecan (ACAN), and collagen 2a1 (COL2A1) mRNA (**Fig. 5**). When loading was initiated immediately (no chondrogenic priming), dynamic loading significantly suppressed SOX9 (**Fig. 5A**) and had no effect on aggrecan (ACAN) expression (**Fig. 5B**), but significantly increased COL2A1 expression (**Fig. 5C**). At priming times of two weeks or more, SOX9 expression was not significantly altered by loading, while both ACAN and COL2A1 expression was significantly increased by dynamic compression. Together, these data indicate a significant effect of duration of priming on mechanotransduction, particularly the response of non-committed hMSCs vs. chondrocytes.

**Figure 5:**
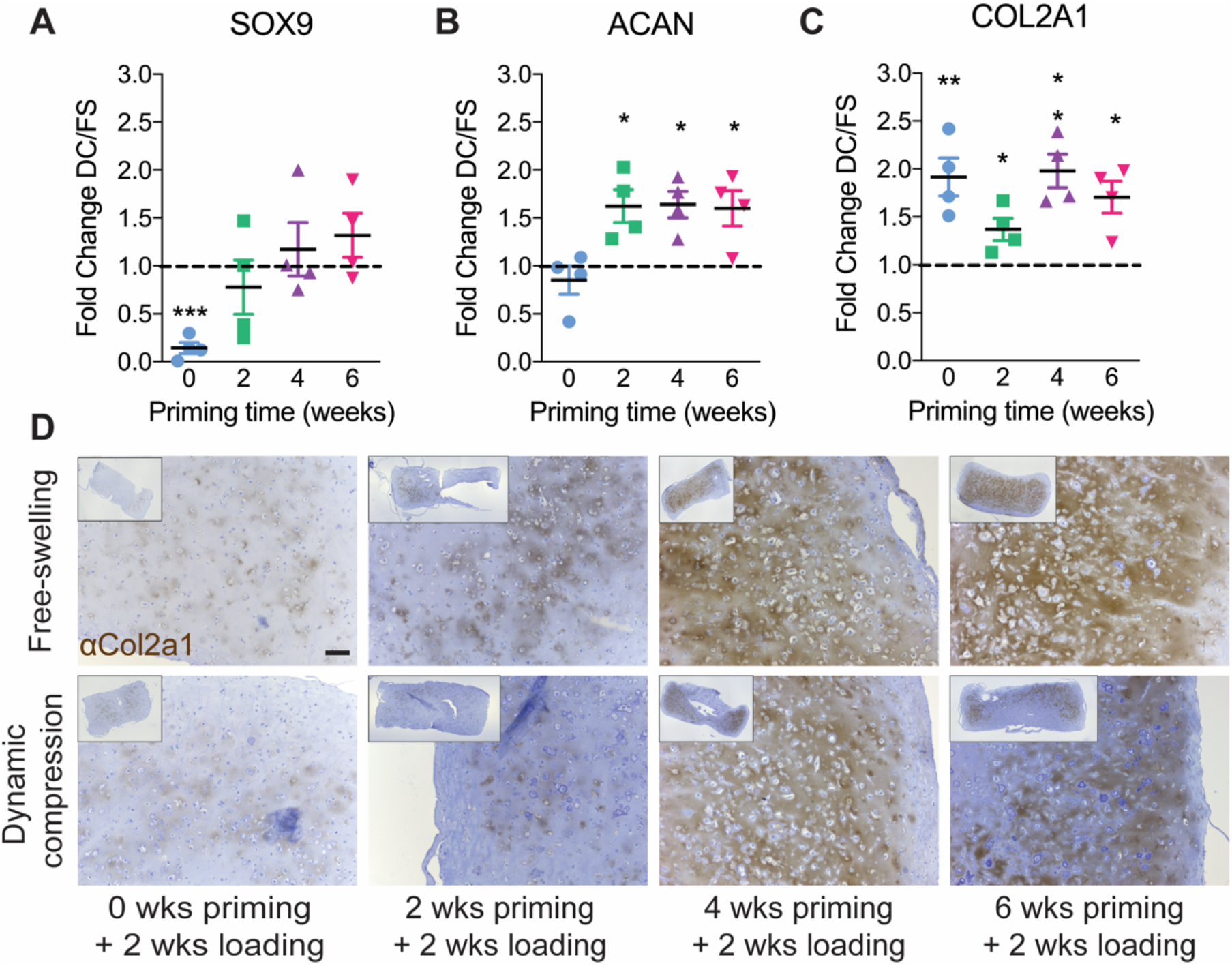
Chondrogenic gene expression under free-swelling (FS) or 10% dynamic compression (DC) conditions. Samples were analyzed immediately after the final loading cycle in each group for mRNA expression of SOX9 (A), ACAN (B), and COL2A1 (C), calculated as fold change over free swelling controls, normalized to GAPDH. Data are shown as mean ± s.d. with individual data points. ANOVA with Tukey’s post-hoc tests were used to determine significance (* p < 0.05, ** p < 0.01). Immunohistochemistry for Col2a1 (D) with hematoxylin counterstain (scale bar = 100 μm). N = 4 per group.

To evaluate the effects of loading on chondrogenic matrix deposition, we performed immunohistochemistry (IHC) staining for Col2a1 protein (**Fig. 5D**). At 0 and 2 weeks priming, Col2a1 staining was minimal regardless of loading conditions. In free-swelling constructs after 4 and 6 weeks of priming, Col2a1 staining was largely more prominent in the construct core compared to dynamic compression.

### Hypertrophic gene expression

Next, to evaluate mechanoregulation of hypertrophic gene expression, we quantified collagen 10a1 (COL10A1) and vascular endothelial growth factor (VEGF) mRNA, which are expressed at high levels by hypertrophic chondrocytes (**Fig. 6**). Dynamic compression significantly increased COL10A1 expression when loading was initiated immediately (no chondrogenic priming) and after 6 weeks of priming (**Fig. 6A**). VEGF regulation was variable with significant upregulation or downregulation depending on priming time (**Fig. 6B**).

**Figure 6:**
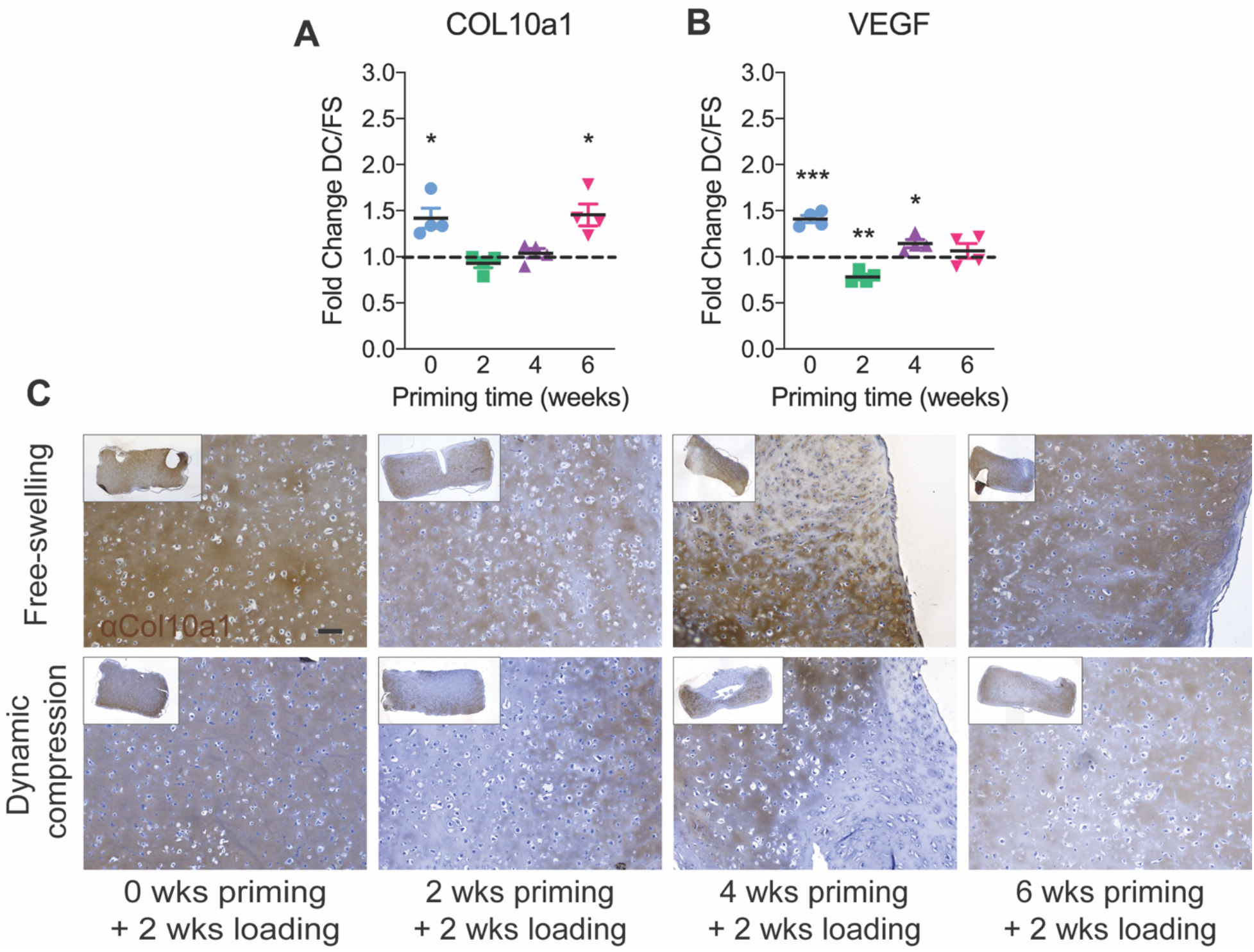
Hypertrophic gene expression under free-swelling (FS) or 10% dynamic compression (DC) conditions. Samples were analyzed immediately after the final loading cycle in each group for mRNA expression of COL10A1 (A) and VEGF (B), calculated as fold change over free swelling controls, normalized to GAPDH. Data are shown as mean ± s.d. with individual data points. ANOVA with Tukey’s post-hoc tests were used to determine significance (* p < 0.05, ** p < 0.01). Immunohistochemistry for Col10a1 (C) with hematoxylin counterstain (scale bar = 100 μm). N = 4 per group.

To evaluate the effects of loading on hypertrophic matrix deposition, we performed immunohistochemistry (IHC) staining for Col10a1 protein (**Fig. 6C**). Under all priming conditions, dynamic compression conditions exhibited lower Col10a1 staining. Similar to Col2a1 distribution, Col10a1 localized throughout the construct core under free-swelling conditions, but exhibited minimal staining at the center under dynamic compression conditions, particularly with increased priming time.

### Osteogenic gene expression

Finally, to evaluate mechanoregulation of osteogenic gene expression, we quantified Runt-related transcription factor 2 (RUNX2) and osteopontin (OPN) mRNA, markers of early and late osteoblastogenesis, respectively. Mechanoregulation of osteogenesis is mediated by the mechanosensitive transcriptional regulator, YAP [35], [36]. We therefore also measured mRNA expression of YAP and the YAP-target gene (through co-activation of the TEAD transcription factors), Cysteine-rich angiogenic inducer 61 (CYR61) (**Fig. 7**) [37]. Loading decreased expression of RUNX2 after 4 weeks of priming, and OPN after 2 and 4 weeks of priming, but the suppressive effect of loading disappeared after 6 weeks of priming (**Fig. 7A,B**). To determine whether this was associated with mechanoregulation of the osteogenic transcriptional regulator, YAP, we evaluated CYR61 as a readout of YAP-TEAD signaling. YAP mechanotransduction is regulated at the protein level, mediated by YAP subcellular localization to the cytosol or the nucleus. As expected, therefore, we found no differences in YAP mRNA expression; however, CYR61 expression was significantly suppressed by dynamic compression after 0, 2, and 4 weeks of chondrogenic priming, in a manner consistent with the induction of chondrogenic gene expression and suppression of osteogenic gene expression (**Fig. 7C,D**).

**Figure 7:**
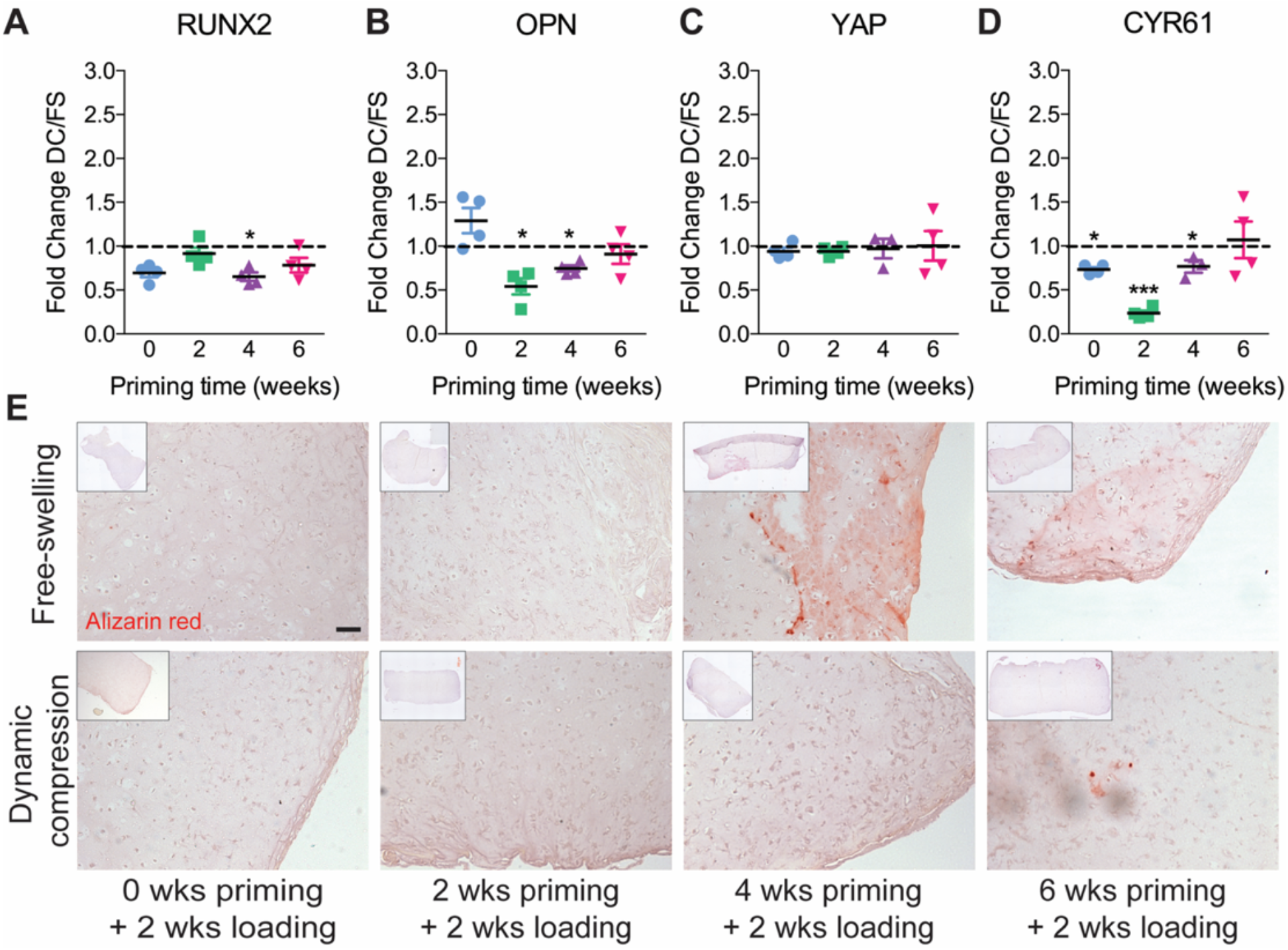
Osteogenic gene expression under free-swelling (FS) or 10% dynamic compression (DC) conditions. Samples were analyzed immediately after the final loading cycle in each group for mRNA expression of RUNX2 (A), osteopontin (OPN) (B), Yes-associated protein (YAP) (C), and the YAP-TEAD transcriptional target gene, Cysteine-rich angiogenic inducer 61 (CYR61) (D), calculated as fold change over free swelling controls, normalized to GAPDH. Data are shown as mean ± s.d. with individual data points. ANOVA with Tukey’s post-hoc tests were used to determine significance (* p < 0.05, ** p < 0.01). Mineral deposition was evaluated by alizarin red staining on histological sections (E) (scale bar = 100 μm). N = 4 per group.

To evaluate the functional effects of loading on mineral deposition, we performed alizarin red staining (**Fig. 7C**). Alizarin red staining was negative in all groups until 4 weeks priming, at which time limited positive staining was observed in free swelling controls but not in those exposed to dynamic compression. After 6 weeks priming, both free-swelling and dynamic compression conditions exhibited modest alizarin red staining, comparable to free-swelling controls after 4 weeks of priming.

## Discussion

In this study, we evaluated the effects of chondrogenic priming duration on the response of engineered human cartilage anlagen to dynamic mechanical compression. We found that the response to dynamic loading depended on the extent of chondrogenic maturation in several ways. First, dynamic loading induced engineered anlage stiffening, in both elastic and viscoelastic behavior, in a manner dependent on the duration of chondrogenic priming prior to load initiation. Further, dynamic loading induced priming time-dependent cellular mechanotransduction and transient expression of extracellular matrix genes, measured by gene expression immediately after the final loading bout. Notably, dynamic loading suppressed type X collagen expression and mineral deposition at latter stages of differentiation. However, the amount and composition of the cartilaginous extracellular matrix depended only on the duration of chondrogenic culture and was not altered by mechanical loading at any time point. Taken together, these observations suggest several hypotheses for future study that may explain the load-increased mechanical properties without changes in bulk matrix amount or composition.

Cartilage mechanical properties are determined by the matrix organization and distribution as well as amount and composition. Thus, mechanical load-induced changes in cartilage matrix organization or spatial distribution may be responsible for the altered mechanical properties. Previous studies on the mechanoresponsiveness of chondrogenically-primed MSCs or primary chondrocytes are consistent with our findings that dynamic loading enhances the mechanical properties of engineered cartilage. For example, studies from Mauck and colleagues, using dynamic loading applied continuously over 9-10 weeks of chondrogenic culture, found that loading increased engineered cartilage mechanical properties by both promoting both type II collagen and GAG deposition [38] and by altering its spatial distribution [39].

In this study, we evaluated matrix deposition after two weeks of loading for different priming times prior to load initiation. Together with our observation that loading transiently increased ECM-regulated gene expression, this suggests that the two-week loading duration may be insufficient to observe substantial changes in matrix composition. However, this load duration was sufficient to alter mechanical properties. We and others have also adopted delayed loading regimes [27], [29], [40], [41] to assess the response of established engineered cartilage to dynamic compression. Similar to the present data, these studies similarly observed increased mechanical properties after delayed loading [29], [40], attributing it mainly to matrix organization [27], [40] or matrix maturation and stress distribution [29]. Despite the short loading duration in this study, we also observed an effect of loading on the spatial distribution of matrix deposition in the constructs. Here, we found that Col2a1 deposition was largely isolated to the core of the hydrogels in the 4- and 6-week priming groups, compared to more uniform distribution under free-swelling conditions. In contrast, Col10a1 was predominantly localized in the periphery of the dynamically loaded hydrogels while free-swelling controls exhibited a more even distribution. These observations are consistent with our prior findings, which suggest that load-induced nutrient transport and/or ECM protein release into the culture bath may contribute to these spatial heterogeneities [27], [40]. Huang et al. similarly observed no effect of loading on average matrix content, but using Fourier-transform infrared spectroscopy (FTIR) found that loading altered the spatial distribution of collagen deposition [29]. Thus, mechanical loading may alter construct mechanical properties, in a manner dependent on the duration of chondrogenic priming by altering extracellular matrix organization and distribution.

Cartilage is a tissue of high cell density, and the mechanical properties of the cells themselves may also contribute significantly to overall tissue mechanical properties. Recently, we discovered a mechanotransductive feedback loop, mediated by the transcriptional regulators YAP and TAZ, that regulates the mechanical properties of the cell itself by modulating cytoskeletal maturation [37]. To maintain cytoskeletal equilibrium, mechano-activated YAP and TAZ drive a transcriptional program that exerts negative feedback on myosin phosphorylation, modulating cytoskeletal tension, and suppression of YAP/TAZ expression or activity increases cell mechanical properties. In this study, we observed that increasing chondrogenic differentiation had no effect on YAP expression, but decreased YAP activity, with further YAP suppression by mechanical loading. This is consistent with our prior findings [5] and with the current literature on the negative role of YAP in chondrocytes [48]. Thus, we speculate that with increased chondrogenic maturation, load-induced suppression of YAP transcriptional activity (measured here by the YAP/TAZ-TEAD target gene, Cyr61) may play a role in the maintenance of a chondrogenic phenotype. With extended maturation (6 weeks of chondrogenic priming), loading no longer suppresses YAP activity, allowing endochondral maturation in the presence of loading. Taken together, we have provided new evidence that dynamic mechanical compression influences engineered cartilage anlage formation in a manner dependent on the chondrogenic maturation state at the time of load initiation. Future research is warranted to dissect the cellular and molecular mechanisms by which chondrocytes receive and interpret mechanical cues and the functional consequences on endochondral bone formation during bone development, repair, and regeneration.

## Limitations

The goal of this study was to observe the effects of chondrogenic priming of human MSCs under dynamic compression, but the study has several limitations that influence our interpretation. First, although MSCs are of potential translational interest for tissue engineering, they may not reflect the biology of the limb bud progenitor cells that give rise to the cartilage anlage in development [36] or of the periosteal progenitor cells that give rise to the fracture callus [49]. Second, MSCs exhibit extensive donor-donor variability. In this study, we used commercially-sourced MSCs from pooled donors, and was not designed to dissect the contributions of donor variability to the observed responses. Third, both gene expression and mechanical testing analyses were performed on bulk tissues, which may mask spatial heterogeneity of cellular mechanotransduction, matrix-associated gene expression, and local mechanical properties. Fourth, this study used only a two-week loading duration. Both shorter and longer durations of mechanical loading would enable dissection of the effects of load timing on acute mechanotransduction and tissue production, respectively. Finally, while our observations of the effects of loading on spatial heterogeneity in matrix deposition are consistent with prior reports from both our own and other groups, low sample sizes precluded histomorphometric quantitation of these outcomes.

## Acknowledgements

This work was supported by the Naughton Foundation (to A.M.M.), the NIH’s National Institute of Arthritis and Musculoskeletal and Skin Diseases under award number R01AR074948 (to JDB), the Penn Center for Musculoskeletal Disorders under award number P30AR069619 (to JDB), and the Center for Engineering Mechanobiology (CEMB), an NSF Science and Technology Center, under grant agreement CMMI: 15-48571. The contents of this publication are solely the responsibility of the authors and do not necessarily represent the official views of the NIH, the National Science Foundation, or other funding agencies.

## Author contributions

D.J.K. and J.D.B. conceived and supervised the research. A.M.M., D.J.K., and J.D.B. designed the experiments. A.M.M., E.A.E., D.J.K., and J.D.B. analyzed the data, and wrote the paper. All authors commented on and approved the final manuscript.

## Notes

### Competing Interest Statement

The authors have declared no competing interest.

## References

[1] S. Provot, E. Schipani, J. Y. Wu, and H. Kronenberg, “Development of the Skeleton,” in Osteoporosis, Elsevier, 2013, pp. 97–126.

[2] C. S. Bahney et al., “Cellular biology of fracture healing,” Journal of Orthopaedic Research, vol. 37, no. 1. John Wiley and Sons Inc., pp. 35–50, Jan-2019.

[3] E. J. Sheehy, D. J. Kelly, and F. J. O’Brien, “Biomaterial-based endochondral bone regeneration: a shift from traditional tissue engineering paradigms to developmentally inspired strategies,” Mater. Today Bio, vol. 3, no. April, p. 100009, 2019.

[4] P. Occhetta et al., “Developmentally inspired programming of adult human mesenchymal stromal cells toward stable chondrogenesis,” Proc. Natl. Acad. Sci. U. S. A., vol. 115, no. 18, pp. 4625–4630, 2018.

[5] A. M. McDermott et al., “Recapitulating bone development through engineered mesenchymal condensations and mechanical cues for tissue regeneration,” Sci. Transl. Med., vol. 11, no. 495, p. eaav7756, Jun. 2019.

[6] E. M. Thompson, A. Matsiko, E. Farrell, D. J. Kelly, and F. J. O’Brien, “Recapitulating endochondral ossification: a promising route to in vivo bone regeneration,” J. Tissue Eng. Regen. Med., vol. 9, no. 8, pp. 889–902, Aug. 2015.

[7] E. M. Thompson, A. Matsiko, D. J. Kelly, J. P. Gleeson, and F. J. O’Brien, “An Endochondral Ossification-Based Approach to Bone Repair: Chondrogenically Primed Mesenchymal Stem Cell-Laden Scaffolds Support Greater Repair of Critical-Sized Cranial Defects Than Osteogenically Stimulated Constructs in Vivo,” Tissue Eng. - Part A, vol. 22, no. 5-6, pp. 556–567, 2016.

[8] A. Matsiko et al., “An endochondral ossification approach to early stage bone repair: Use of tissue-engineered hypertrophic cartilage constructs as primordial templates for weight-bearing bone repair,” J. Tissue Eng. Regen. Med., vol. 12, no. 4, pp. e2147–e2150, 2018.

[9] E. Alsberg, K. W. Anderson, A. Albeiruti, J. A. Rowley, and D. J. Mooney, “Engineering growing tissues,” Proc. Natl. Acad. Sci. U. S. A., vol. 99, no. 19, pp. 12025–12030, Sep. 2002.

[10] W. Roux, Gesammelte Abhandlungen über Entwickelungsmechanik der Organismen: Bd. Entwicklungsmechanik des Embryo. Wilhelm Engelmann, 1895.

[11] V. Hamburger and M. Waugh, “The Primary Development of the Skeleton in Nerveless and Poorly Innervated Limb Transplants of Chick Embryos,” Physiol. Zool., vol. 13, no. 4, pp. 367–382, 1940.

[12] N. C. Nowlan, J. Sharpe, K. A. Roddy, P. J. Prendergast, and P. Murphy, “Mechanobiology of embryonic skeletal development: Insights from animal models,” Birth Defects Res. Part C Embryo Today Rev., vol. 90, no. 3, pp. 203–213, Sep. 2010.

[13] S. W. Verbruggen et al., “Stresses and strains on the human fetal skeleton during development.,” J. R. Soc. Interface, vol. 15, no. 138, p. 20170593, Jan. 2018.

[14] N. Felsenthal and E. Zelzer, “Mechanical regulation of musculoskeletal system development,” Development, vol. 144, no. 23,pp. 4271–4283, Dec. 2017.

[15] R. Hente, B. Füchtmeier, U. Schlegel, a Ernstberger, and S. M. Perren, “The influence of cyclic compression and distraction on the healing of experimental tibial fractures.,” J. Orthop. Res., vol. 22, no. 4, pp. 709–15, 2004.

[16] S. M. Perren and J. Cordey, “The concept of interfragmentary strain,” Curr. concepts Intern. Fixat. Fract., pp. 63–77, 1980.

[17] A. E. Goodship, P. E. Watkins, H. S. Rigby, and J. Kenwright, “The role of fixator frame stiffness in the control of fracture healing. An experimental study,” J. Biomech., vol. 26, no. 9, pp. 1027–1035, 1993.

[18] L. E. Claes et al., “Effects of mechanical factors on the fracture healing process,” Clin. Orthop. Relat. Res., no. 355 SUPPL., 1998.

[19] P. Klein et al., “The initial phase of fracture healing is specifically sensitive to mechanical conditions,” J. Orthop. Res., vol. 21, no. 4, pp. 662–669, 2003.

[20] R. L. Duncan and C. H. Turner, “Mechanotransduction and the Functional Response of Bone to Mechanical Strain R.,” Calcif. Tissue Int., vol. 57, pp. 344–358, 1995.

[21] L. E. Claes and C. A. Heigele, “Magnitudes of local stress and strain along bony surfaces predict the course and type of fracture healing,” J. Biomech., vol. 32, no. 3, pp. 255–266, 1999.

[22] L. Claes, A. Veeser, M. Göckelmann, U. Simon, and A. Ignatius, “A novel model to study metaphyseal bone healing under defined biomechanical conditions,” Arch. Orthop. Trauma Surg., vol. 129, no. 7, pp. 923–928, 2009.

[23] L. Claes, S. Recknagel, and A. Ignatius, “Fracture healing under healthy and inflammatory conditions.,” Nat. Rev. Rheumatol., vol. 8, no. 3, pp. 133–43, Jan. 2012.

[24] S. Herberg et al., “Combinatorial morphogenetic and mechanical cues to mimic bone development for defect repair,” Sci. Adv., vol. 5, no. 8, p. eaax2476, Aug. 2019.

[25] J. D. Boerckel, B. a Uhrig, N. J. Willett, N. Huebsch, and R. E. Guldberg, “Mechanical regulation of vascular growth and tissue regeneration in vivo.,” Proc. Natl. Acad. Sci. U. S. A., vol. 108, no. 37, pp. E674–80, Aug. 2011.

[26] M. A. Ruehle et al., “Extracellular matrix compression temporally regulates microvascular angiogenesis,” Sci. Adv., vol. 6, no. 34, p. eabb6351, Aug. 2020.

[27] S. D. Thorpe, C. T. Buckley, T. Vinardell, F. J. O’Brien, V. A. Campbell, and D. J. Kelly, “The response of bone marrow-derived mesenchymal stem cells to dynamic compression following tgf-β3 induced chondrogenic differentiation,” Ann. Biomed. Eng., vol. 38, no. 9, pp. 2896–2909, 2010.

[28] M. G. Haugh, E. G. Meyer, S. D. Thorpe, T. Vinardell, G. P. Duffy, and D. J. Kelly, “Temporal and spatial changes in cartilage-matrix-specific gene expression in mesenchymal stem cells in response to dynamic compression,” Tissue Eng. - Part A, vol. 17, no. 23-24, pp. 3085–3093, 2011.

[29] A. H. Huang, M. J. Farrell, M. Kim, and R. L. Mauck, “LONG-TERM DYNAMIC LOADING IMPROVES THE MECHANICAL PROPERTIES OF CHONDROGENIC MESENCHYMAL STEM CELL-LADEN HYDROGELS,” Eur. Cells Mater., vol. 19, pp. 72–85, 2010.

[30] T. Zhang et al., “Biomaterials Cross-talk between TGF-beta / SMAD and integrin signaling pathways in regulating hypertrophy of mesenchymal stem cell chondrogenesis under deferral dynamic compression,” Biomaterials, vol. 38, pp. 72–85, 2015.

[31] Y. J. Kim, R. L. Y. Sah, J. Y. H. Doong, and A. J. Grodzinsky, “Fluorometric assay of DNA in cartilage explants using Hoechst 33258,” Anal. Biochem., vol. 174, no. 1, pp. 168–176, Oct. 1988.

[32] W. Kafienah and T. J. Sims, “Biochemical methods for the analysis of tissue-engineered cartilage.,” Methods Mol. Biol., vol. 238, pp. 217–230, 2004.

[33] N. Y. Ignat’eva, N. A. Danilov, S. V. Averkiev, M. V. Obrezkova, V. V. Lunin, and E. N. Sobol’, “Determination of hydroxyproline in tissues and the evaluation of the collagen content of the tissues,” J. Anal. Chem., vol. 62, no. 1, pp. 51–57, Jan. 2007.

[34] T. D. Schmittgen and K. J. Livak, “Analyzing real-time PCR data by the comparative CT method,” Nat. Protoc., vol. 3, no. 6, pp. 1101–1108, Jun. 2008.

[35] S. Dupont et al., “Role of YAP/TAZ in mechanotransduction.,” Nature, vol. 474, no. 7350, pp. 179–83, Jun. 2011.

[36] C. D. Kegelman et al., “Skeletal cell YAP and TAZ combinatorially promote bone development,” FASEB J., p. fj.201700872R, Jan. 2018.

[37] D. E. Mason et al., “YAP and TAZ limit cytoskeletal and focal adhesion maturation to enable persistent cell motility,” J. Cell Biol., vol. 218, no. 4, pp. 1369–1389, Apr. 2019.

[38] R. L. Mauck, S. L. Seyhan, G. A. Ateshian, and C. T. Hung, “Influence of seeding density and dynamic deformational loading on the developing structure/function relationships of chondrocyte-seeded agarose hydrogels,” Ann. Biomed. Eng., vol. 30, no. 8, pp. 1046–1056, 2002.

[39] L. Bian et al., “Dynamic mechanical loading enhances functional properties of tissue-engineered cartilage using mature canine chondrocytes,” Tissue Eng. - Part A, vol. 16, no. 5, pp. 1781–1790, 2010.

[40] L. Luo, S. D. Thorpe, C. T. Buckley, and D. J. Kelly, “The effects of dynamic compression on the development of cartilage grafts engineered using bone marrow and infrapatellar fat pad derived stem cells,” Biomed. Mater., vol. 10, no. 5, 2015.

[41] L. Bian, D. Y. Zhai, E. C. Zhang, R. L. Mauck, and J. A. Burdick, “Dynamic compressive loading enhances cartilage matrix synthesis and distribution and suppresses hypertrophy in hMSC-laden hyaluronic acid hydrogels.,” Tissue Eng. Part A, vol. 18, no. 7-8, pp. 715–24, 2012.

[42] M. J. Grimshaw and R. M. Mason, “Bovine articular chondrocyte function in vitro depends upon oxygen tension,” Osteoarthr. Cartil., vol. 8, no. 5, pp. 386–392, 2000.

[43] C. T. Buckley, E. G. Meyer, and D. J. Kelly, “The influence of construct scale on the composition and functional properties of cartilaginous tissues engineered using bone marrow-derived mesenchymal stem cells,” Tissue Eng. - Part A, vol. 18, no. 3-4, pp. 382–396, 2012.

[44] M. C. Lewis, B. D. MacArthur, J. Malda, G. Pettet, and C. P. Please, “Heterogeneous proliferation within engineered cartilaginous tissue: The role of oxygen tension,” Biotechnol. Bioeng., vol. 91, no. 5, pp. 607–615, 2005.

[45] F. Witt, G. N. Duda, C. Bergmann, and A. Petersen, “Cyclic mechanical loading enables solute transport and oxygen supply in bone healing: An in vitro investigation,” Tissue Eng. - Part A, vol. 20, no. 3-4, pp. 486–493, 2014.

[46] L. Bian et al., “Influence of decreasing nutrient path length on the development of engineered cartilage,” Osteoarthr. Cartil., vol. 17, no. 5, pp. 677–685, May 2009.

[47] A. C. Daly, B. N. Sathy, and D. J. Kelly, “Engineering large cartilage tissues using dynamic bioreactor culture at defined oxygen conditions,” J. Tissue Eng., vol. 9, 2018.

[48] A. Karystinou, A. J. Roelofs, A. Neve, F. P. Cantatore, H. Wackerhage, and C. De Bari, “Yes-associated protein (YAP) is a negative regulator of chondrogenesis in mesenchymal stem cells,” Arthritis Res. Ther., vol. 17, no. 1, pp. 1–14, 2015.

[49] C. D. Kegelman et al., “YAP and TAZ promote periosteal osteoblast precursor expansion and differentiation for fracture repair,” J. Bone Miner. Res., vol. in press, p. 2020.03.17.995761, Mar. 2020.

